# Motor properties of PilT-independent type 4 pilus retraction in gonococci

**DOI:** 10.1101/497933

**Authors:** Robert Zöllner, Tom Cronenberg, Berenike Maier

**Affiliations:** University of Cologne, Institute for Biological Physics

**Keywords:** Pilus, molecular motor, twitching motility, Neisseria gonorrhoeae

## Abstract

Bacterial type 4 pili (T4P) belong to the strongest molecular machines. The gonococcal T4P retraction ATPase PilT supports forces exceeding 100 pN during T4P retraction. Here, we address the question whether gonococcal T4P retract in the absence of PilT. We show that *pilT* deletion strains indeed retract their T4P but the maximum force is reduced to 5 pN. Similarly, the speed of T4P retraction is lower by orders of magnitude compared to T4P retraction driven by PilT. Deleting the *pilT* paralogues *pilU* and *pilT2* in the *ΔpilT* background did not inhibit T4P retraction, indicating that the PilT-like proteins do not compensate for PilT. Furthermore, we show that depletion of proton motive force did not inhibit pilT-independent T4P retraction. We conclude that the retraction ATPase is not essential for gonococcal T4P retraction. However, the force generated in the absence of PilT is too low to support important functions of T4P including twitching motility, fluidization of colonies, or induction of host cell response.

**IMPORTANCE:** Bacterial type 4 pili (T4P) have been termed the ‘swiss army knive’ of bacteria because they perform numerous functions including host cell interaction, twitching motility, colony formation, DNA uptake, protein secretion, and surface sensing. The pilus fibre continuously elongates or retracts and these dynamics are functionally important. Curiously, only a subset of T4P systems employs T4P retraction ATPases to power T4P retraction. Here we show that one of the strongest T4P machines, the gonococcal T4P, retracts without a retraction ATPase. Biophysical characterization reveal strongly reduced force and speed compared to retraction with ATPase. We propose that bacteria encode for retraction ATPases when T4P have to generate high force supporting functions like twitching motility, triggering host cell response, or fluidizing colonies.

## INTRODUCTION

Bacterial type 4 pili (T4P) are among the strongest molecular machines known to date. In some species they generate forces exceeding 100 pN (1–3), i.e. twenty-fold higher than the force generated by muscle myosin. Force generation has been linked to diverse functions including twitching motility (4–7), host cell interaction (8–11), and regulation of biofilm structure and dynamics (12–17). For all of these functions, the retraction ATPase PilT is required. Interestingly, some T4P systems involved in protein secretion, DNA uptake during transformation, or surface sensing bear no *pilT*-like gene. Very recently, it has been shown that T4P can retract in the absence of a retraction ATPase (18–20). The forces generated by these pili, however, are by an order of magnitude lower than the force observed for *Neisseria gonorrhoeae* T4P retraction. It remains unclear, whether gonococci can retract T4P in the absence of PilT.

The T4P filament is a helical structure built from thousands of major pilin subunits and various minor pilins (21, 22). Recent advances in cryo-electron microscopy together with high-resolution structures of the individual components have given insight into the structure of the complex machinery that shuttles pilin subunits from the cytoplasmic membrane into the growing pilus (23–27) (Fig. 1). The motor subcomplex comprises the cytoplasmic ATPases that power elongation (PilF) (28) and retraction (PilT) (29) of the pilus, respectively, and an inner membrane platform protein (PilG). The outer membrane subcomplex formed by PilQ enables T4P extrusion (30). The alignment subcomplex spans both outer and the cytoplasmic membrane connecting motor and secretin subcomplexes (23). The T4P elongation and retraction ATPases belong to the family of secretion ATPases. Biophysical and electron microscopy studies suggest that these motors form hexameric rings (31, 32). Interestingly, the structures show oblong, two-fold symmetric hexamers. Each pair of opposing subunits adopts a nucleotide-dependent state with different spatial arrangements (32). Based on crystal structures with different nucleotides it was proposed that sequential ATP binding leads to functionally relevant deformations that propagate around the ring in opposite directions for the elongation and retraction ATPase (24). These sequential deformations of the ATPases would couple to the pilus fibre through the platform complex. In particular, the platform complex was proposed to rotate in response to the conformational changes of the hexameric ATPases (23, 24). Since the T4P fibre is helical, opposite rotations driven by PilF and PilT would then power elongation and retraction of the T4P fibre, respectively. The exact coupling mechanism remains unclear.

**FIG 1.**
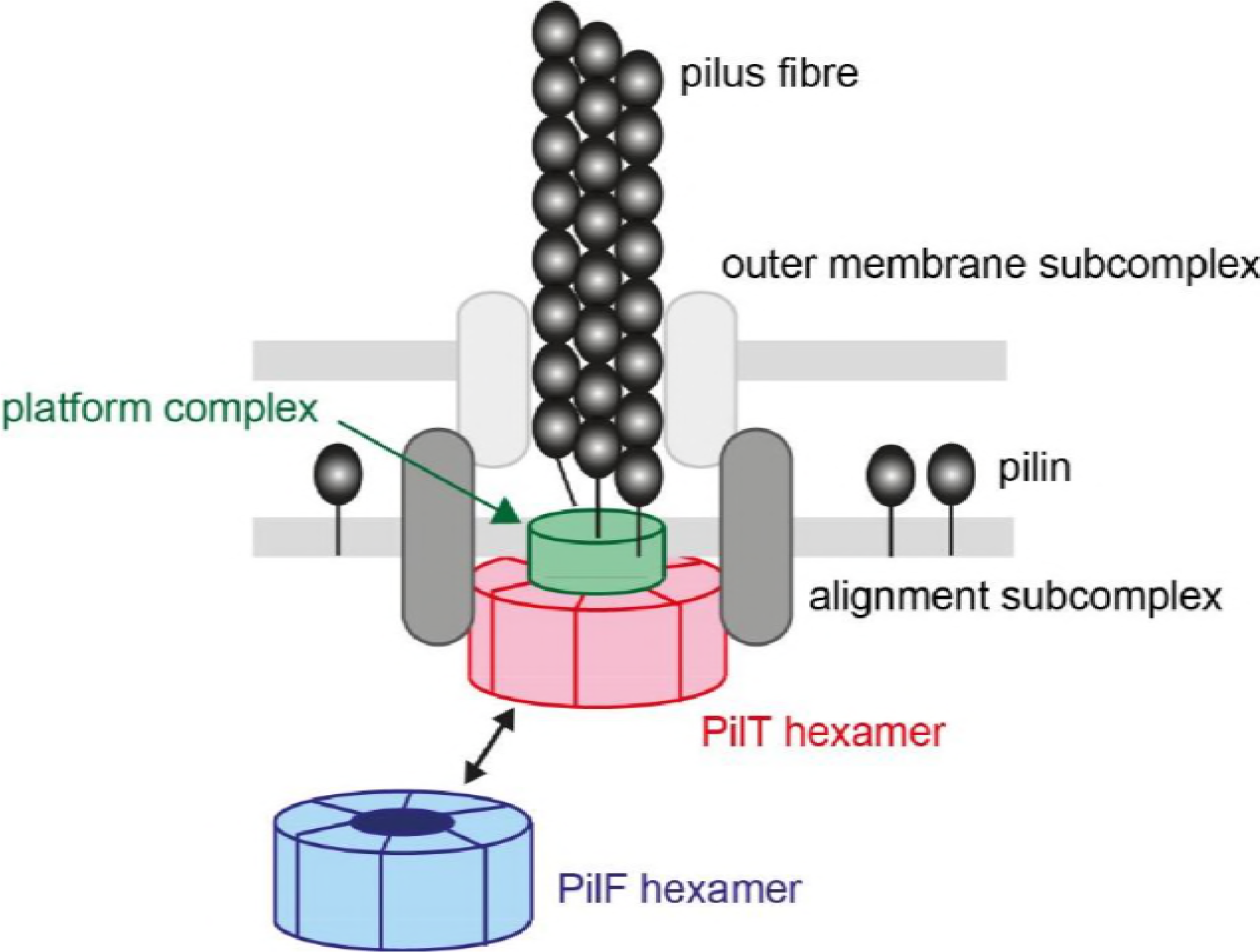
Hypothetical model of the T4P machine. The outer membrane subcomplex enables secretion of the pilus fibre. The alignment subcomplex is in contact with the outer membrane subcomplex and the motor subcomplex comprising the elongation ATPase PilF, the retraction ATPase PilT, and the platform complex formed by PilG. PilF and PilT form oblong, two-fold symmetric hexamers. Sequential ATP binding and hydrolysis causes a deformation wave running through the rings in opposite directions for PilF and PilT hexamers. This deformation wave couples to the platform complex, potentially causing platform rotation. Due to the helical shape of the pilus fibre, the direction of rotation determines whether pilins are inserted into or removed from the terminal end of the fibre.

While all bacteria generating T4P encode for elongation ATPases, not all encode for retraction ATPases. For example, DNA uptake during transformation has been reported to require a retraction ATPase in *N. gonorrhoeae* and *V. cholerae* (29, 33). However, *B. subtilis* and S. *pneumoniae* do not carry a clear *pilT* homologue, but still they employ T4P for DNA uptake (34). The first T4P system shown to retract T4P in the absence of a retraction ATPase was the toxin co-regulated pilus of *V. cholerae* (18). The maximum force generated by these T4P was in the range of 4 pN. Furthermore, the Tad pilus of *Caulobacter crescentus* generates somewhat higher force in the range of 12 pN (19). We note that it is unknown how force generation depends on experimental conditions. For two T4P systems that naturally encode for a retraction ATPase, force generation was observed when *pilT* was deleted. First, in *Myxococcus xanthus* deletion of *pilT* leads to strong reduction of T4P retraction frequency and almost complete loss of twitching motility (2). The force was reduced two-fold. *M. xanthus* carries four *pilT* paralogues and it is therefore unclear whether any of them encodes for a functional retraction ATPase. Second, the competence pilus of *V. cholerae* showed T4P retraction when *pilT* was deleted (20) while the transformation rate was severely reduced. Forces generated by the competence pilus were on the lower side (8 pN) even in the presence of functional PilT and dropped by two-fold in a *pilT* deletion strain. Therefore, it was interesting to find out how severely deletion of *pilT* and its paralogues affected force generation in *N. gonorrhoeae*, one of the strongest T4P machines.

Here, we set out to address this question by characterizing velocity and force generation in gonococcal *pilT* deletion strains. We show that indeed gonococcal T4P retract independent of *pilT* and its paralogues. Interestingly, both force and speed of PilT-independent T4P retraction are lower by orders of magnitude compared to retraction in wt T4P, explaining why T4P retraction has been overlooked so far. We investigate putative energy sources of PilT-independent T4P retraction by characterizing the effects of proton motive force and pilin concentration.

## RESULTS

### T4P retract at low speed in a *pilT* deletion mutant

Deleting the gene encoding for the T4P retraction ATPase PilT was long believed to be in accord with generating strains that are incapable of T4P retraction (4). However, recent experiments reported T4P retraction in the absence of *pilT* or homologues (20). We set out to investigate whether T4P retraction occurred in gonococcal *pilT* deletion strains. We used a laser tweezers assay to probe for T4P retraction (Fig. 2a). A bacterium was immobilized at a glass coverslide. Using a laser trap, a polystyrene bead was placed adjacent to the bacterium. When a T4P bound to the bead and retracted, the bead was deflected by a distance *d* from the center of the laser trap. The force acting on the bead is proportional to the optical restoring force *F.* In force clamp mode, *d* was kept constant by moving the microscope stage by a distance *δ* with respect to the center of the laser trap. Thus by measuring *δ*, we can determine the velocity of T4P retraction at constant force.

**FIG 2.**
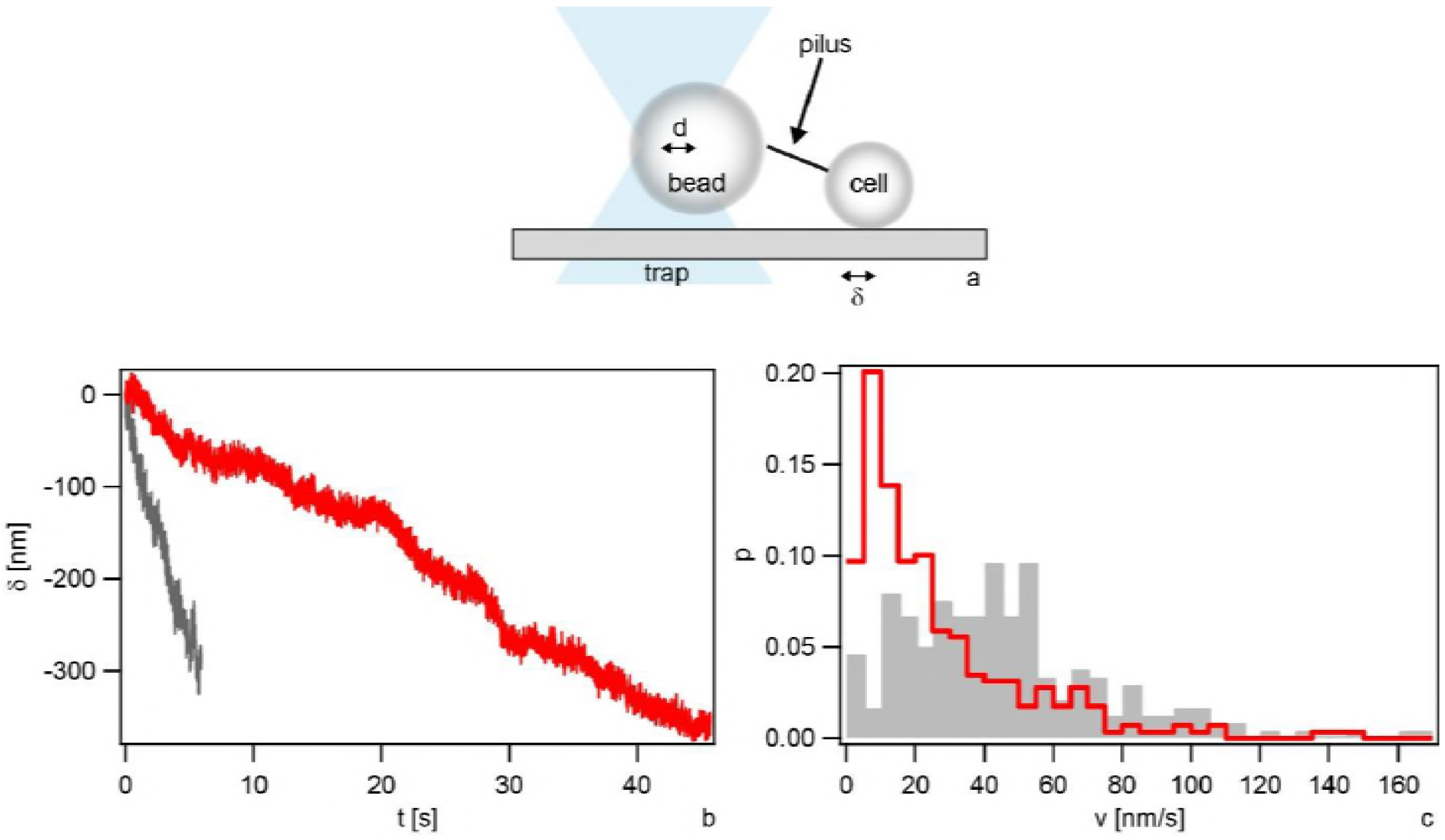
*pilT*-independent T4P retraction in strain *ΔpilT* (Ng178). a) Sketch of the experimental setup. A gonococcal cell is attached to a glass surface and a polystyrene bead trapped in a laser trap is placed in close proximity. When a T4P binds to the bead, retraction deflects the bead from the center of the trap by a distance *d*. *d* is proportional to the optical restoring force *F*. In force clamp mode, *F* is held constant by moving the surface-bound cell by a distance *δ*. Thus *δ* is a measure of the T4P length change. b) Typical examples of T4P length change *δ* as a function of time and c) velocity distribution of at forces clamped at F = 4 pN (grey, N = 239) and F = 8pN (red, N = 298), respectively.

To start with, we clamped the force at F = 8 pN. This is the lowest force that we typically used for characterizing T4P retraction in the wt strain with functional PilT. We found that the *ΔpilT* strain indeed deflected the bead from the center of the laser trap, indicating T4P retraction (Fig. 2b). The distribution of velocities showed a maximum around v = 5 nm s^−1^ and a pronounced tail towards higher velocities (Fig. 2c). We assessed whether similar retractile behavior occurred in a different trapping geometry. A dual trap (17) was used to trap a single spherical bacterium in each trap. This setup was not influenced by microscope drift. Again, T4P retraction was observed (Fig. S1a). Furthermore, we used a configuration where we trapped a spherical bacterium in one trap and a bead in the second trap. Likewise, T4P retraction was observed (Fig. S1b). We conclude that gonococcal T4P retract in the absence of the retraction ATPase PilT.

Next, the effect of force on the speed of PilT-independent T4P retraction was investigated. We measured the velocity when the force was clamped to F = 4 pN (Fig. 2b). As expected, the distribution of velocities shifted to higher values as compared to F = 8 pN. The distribution showed a maximum around v = 40 nm s^−1^ and again a tail at higher velocities (Fig. 2c). No clear T4P elongation events were observed.

In summary, gonococcal T4P retract in the absence of the retraction ATPase PilT, but the speed is by two orders of magnitude lower compared to the speed measured in the PilT-producing strain.

### PilT-independent T4P retraction generates lower force compared to PilT-drivenretraction

Gonococcal T4P are among the strongest molecular machines described so far (5,35). We addressed force generation in the absence of the T4P retraction ATPase PilT. To this end, T4P retraction was probed using the laser trap in the position clamp mode. As the T4P pulled on the bead the deflection increased and concomitantly, the force increased. Eventually, the speed leveled off (Fig. 3a). A stalling event was defined as an event where *dF/dt* = 0 for 1s or longer. The stalling forces were distributed around a mean (± sd) of F = (4.7 ± 0.7) pN (Fig. 3b). We note that there were outliers at considerably higher forces. They were disregarded when calculating the average stalling force. Most likely, these stalling events have been caused by multiple T4P pulling on the bead simultaneously.

**FIG 3.**
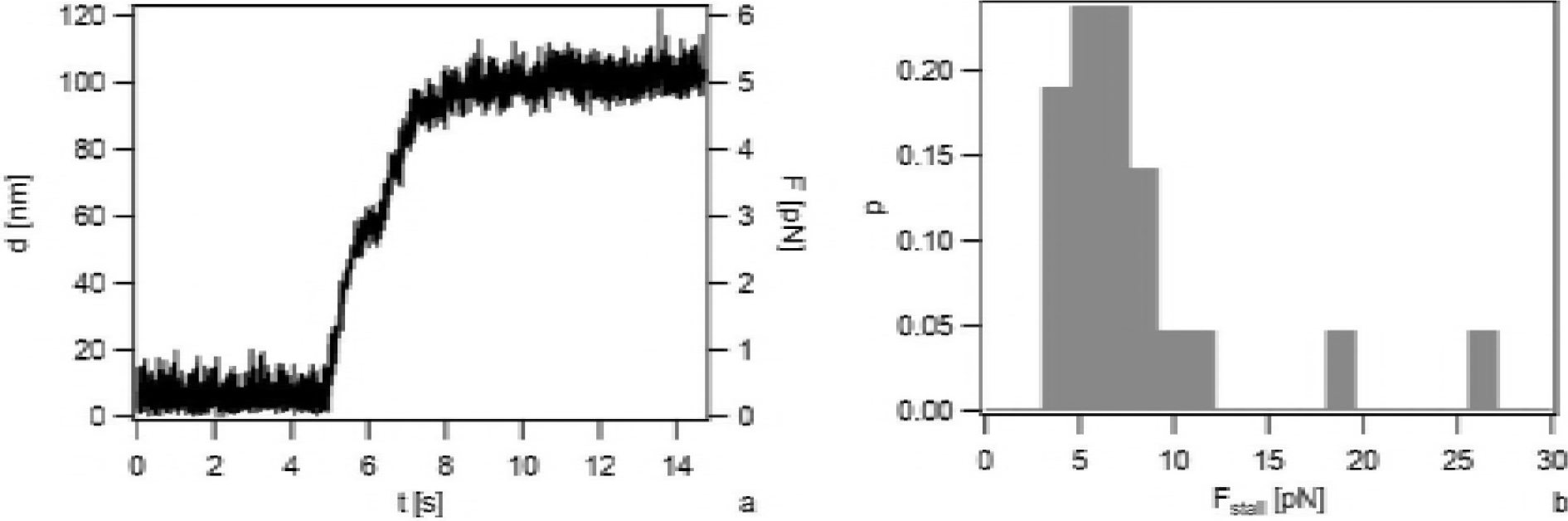
Stalling of *pilT*-independent T4P retraction in strain *ΔpilT* (Ng178). a) Typical stalling event in position clamp mode. Deflection of the bead from the center of the laser trap *d* and force *F* are plotted as function of time. b) Distribution of stalling forces *F_stall_*. N = 21.

We conclude that PilT-independent T4P retraction generates force in the range of 5 pN, i.e. 20-30 fold lower compared to PilT-powered T4P retraction.

### T4P retraction without PilT occurs independently of the *pilT* paralogues *pilU* and *pilT2*

*N. gonorrhoeae* bears two *pilT* paralogues on its genome, namely *pilU* and *pilT2*. *pilU* resides in the same operon as *pilT* and its deletion shows little effect on T4P retraction in wt gonococci (36). Deletion of *pilT2* in the wt background causes a reduction of T4P retraction speed by a factor of ~ 2 (36). It was conceivable that one of the proteins encoded by *pilT* paralogues powered PilT-independent T4P retraction. Using the dual trap assay, we probed to following strains for T4P retraction; *ΔpilT2 ΔpilT, ΔpilU ΔpilT, and ΔpilT2 ΔpilU ΔpilT*. All three strains showed T4P retraction (Table 1) indicating that the two gonococcal paralogues are not responsible for T4P retraction in the absence of PilT.

**TABLE 1.** Deletion of the gonococcal *pilT* paralogues does not inhibit T4P retraction.

| Strain | Genotype | Force Generation | Reference |
| --- | --- | --- | --- |
| $\Delta G4$ (Ng150) | *G4::aac* | yes | (48) |
| $\Delta pilT$ (Ng178) | *pilT::m-Tn3cm*<br>*G4::aac* | yes | (17) |
| $\Delta pilT \Delta pilT2$ (Ng184) | *pilT2::kanR*<br>*pilT::m-Tn3cm*<br>*G4::aac* | yes | (36), this study |
| $\Delta pilT \Delta pilU$ (Ng185) | *pilU::ermC*<br>*pilT::m-Tn3cm*<br>*G4::aac* | yes | (36), this study |
| $\Delta pilT \Delta pilT2 \Delta pilU$ (Ng186) | *pilT2::kanR*<br>*pilU::ermC*<br>*pilT::m-Tn3cm*<br>*G4::aac* | yes | (36), this study |
| *3xpilE* (Ng188) | *igA1::P<sub>pilE</sub>pilE P<sub>pilE</sub> pilE ermC*<br>*pilT::m-Tn3cm*<br>*G4::aac* | yes | (39, 40), this study |

### Depletion of proton motive force does not inhibit PilT-independent T4P retraction

Depletion of proton motive force reduces the speed of gonococcal T4P retraction two-fold (37,38). To test whether proton motive force powers *pilT*-independent T4P retraction, Δ*pilT* cells were treated with the uncoupler carbonyl cyanidem-chlorophenyl hydrazone (CCCP). CCCP shuttles protons across the membrane, in the direction of the proton gradient and deplete PMF. Cells were incubated with 50 μM CCCP for 15 min prior to usage in dual laser tweezers. Notably, Δ*pilT* cells were able to generate forces between 15 min and and 50 min post treatment with CCCP-treatment. We conclude that PilT-independent retraction is not driven by proton motive force.

### The speed of PilT-independent T4P retraction depends on the concentration of the major pilin

Finally, we investigated the effect of overproducing the major pilin PilE. We used strain *3xpilE* that carries two tandemly arrayed (identical) *pilE* genes expressed ectopically in addition to the *pilE* gene in the native locus (39). It was constructed essentially as described for strain GE21 (40) with the exception that it carries two gene copies in addition to the native copy resulting in a strain expressing three identical *pilE* genes. In this strain, additionally *pilT* was deleted. Using the single laser trap in force clamp mode, we measured the velocity of PilT-independent T4P retraction. The velocity distribution in the pilin-overproducing strain was shifted towards lower values compared to the distribution of the strain with only the native *pilE* copy (Fig. 4).

**FIG 4.**
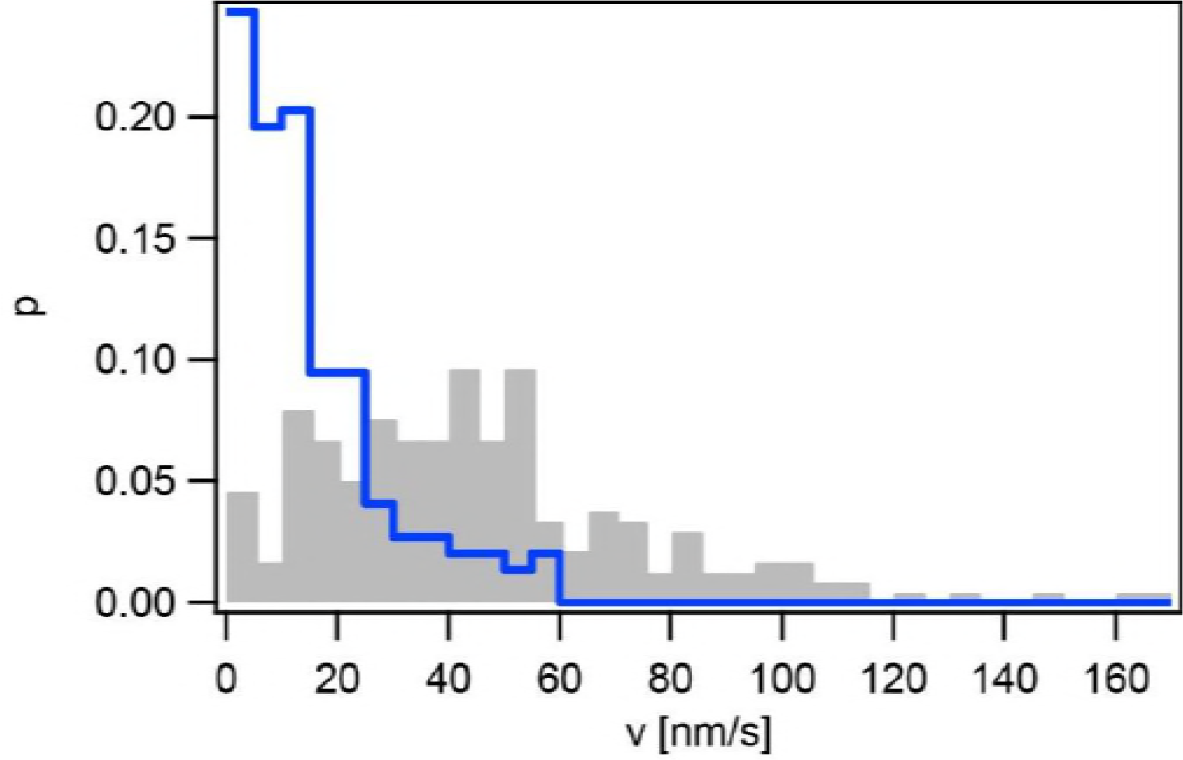
T4P retraction velocity depends on concentration of pilin. Velocity distribution of *ΔpilT* (grey, Ng178, N = 239) and pilin over producing strain P_pilE_*pilE* P_pilE_*pilE ΔpilT* (blue, Ng188, N = 148) at F = 4 pN.

To summarize, over expressing the major pilin *pilE* results in reduced velocity of PilT-independent T4P retraction.

## DISCUSSION

### Comparing T4P retraction in the presence and absence of the T4P retraction ATPase

It is important to note that so far a gonococcal *pilT* deletion strain was considered to incapable of T4P retraction (4, 9, 17, 41, 42). In our own studies characterizing single T4P retraction (1, 35) we have overlooked retraction in the absence of *pilT* because the velocity is close to zero in the force range we have worked so far. Most other studies have probed for T4P retraction by measuring phenotypic consequences of T4P retraction including twitching motility or DNA transformation. Indeed, deleting *pilT* inhibits gonococcal twitching motility and transformation (4, 29, 43).

T4P retraction in the absence of the retraction ATPase PilT has been recently reported for various bacterial species. First, T4P retraction was reported in T4P systems that naturally lack the gene encoding for the T4P retraction ATPase (18, 19). Second, in T4P systems that naturally bear genes encoding for the T4P retraction ATPase, the latter was deleted and still T4P retraction was observed (2, 20). Here, we show that deletion of *pilT* in the gonococcal T4P system has a strong effect on force generation. While wt T4P generate force in the range of 150 pN (5), the stalling forces measured for the *ΔpilT* strain are in the range of 5 pN. Moreover, the retraction velocity is strongly reduced. It is interesting to compare the forces generated by various T4P systems in the presence and absence of the retraction ATPase. In the absence of the retraction ATPase, the forces generated by T4P retraction are fairly low in the range of (3–12) pN (18, 19). The exception is T4P retraction in a *M. xanthus ΔpilT* strain where force in the range of 70 pN are generated (2). However, it is likely that one of the *pilT* paralogues powers retraction for this specific strain. On the other hand, T4P retraction powered by the retraction ATPase shows a wide range of forces from 8 pN up to 150 pN (2, 5, 20, 44). Taken together, we conclude that the retraction ATPase PilT consistently increases the force generated by T4P retraction in different bacterial species.

### Implications of retraction ATPase-independent T4P retraction for chemo-mechanical coupling in the T4P machine

Recently, progress has been made in understanding the chemo-mechanical coupling in the T4P machine (Fig. 1). Most likely, ATP binding and / or hydrolysis induces conformational changes of the retraction ATPase hexamer parallel to the membrane (23, 24, 32). Structural data is consistent with a wave of conformational changes around the ring leading to rotational motion of the platform complex. T4P retraction force, however, is generated perpendicular to the membrane by collapse of pilins from the fibre into the membrane and it remains to be determined how putative rotation of the platform complex shuffles pilins from the fibre into the cytoplasmic membrane. The fact that T4P retraction occurs in the absence of the retraction ATPase evokes speculations about the energetics of the T4P machine. Deleting the gene encoding for the elongation ATPase leads to non-piliated bacteria (28). So far, no T4P lacking the elongation ATPase has been reported to our knowledge. These two facts strongly suggest that the energy provided by ATP hydrolysis in the elongation ATPase is required for T4P polymerization. Part of this energy may be stored in the T4P fibre and power T4P depolymerization when the elongation ATPase has dissociated from the complex. Currently, an interesting open question is how the conformational changes in the platform complex control insertion and removal of the terminal pilins. It is conceivable, that thermal fluctuations are biased towards the direction of removal in the absence of any ATPase. Thus an elongation ATPase is essential for T4P polymerization. The retraction ATPase would not be strictly required for depolymerization, but biasing the conformational change of the platform complex in the direction of pilin removal would speed up the process.

Notably, increasing *pilE* expression reduces the velocity of T4P retraction. Assuming that the concentration of pilin in the cytoplasmic membrane is increased in the PilE-overproducing strain, entropic effects may explain why the velocity is lower. Consider the terminal pilin of the pilus fibre. We assume that a free energy landscape describes transitions of this pilin between the fibre and the membrane. Current data is consistent with the free energy of the membrane-inserted state being lower than the free energy of the pilus-inserted state. Increasing the concentration of pilins in the membrane would then change the free energy landscape favoring the pilus-inserted state. At this point, however, we cannot exclude other causes for reduced velocity including interaction with other pili that may increase friction during T4P retraction. Interestingly, little T4P elongation was observed in this study. If the retraction and elongation ATPases bound alternatively and stochastically, then we would expect to see occasional switching from slow T4P retraction to fast elongation (indicative of binding of the elongation ATPase). The lack of T4P elongation in our study may suggest that T4P have to retract fully prior to binding of the elongation ATPase. This finding is consistent with processive retraction of toxin co-regulated pili in *V. cholerae* where it was proposed that insertion of minor pilins blocked elongation and triggered retraction (18). Previously, we observed that T4P retraction frequently switched to elongation in gonococcal strains that had strongly reduced concentrations of PilT (35, 45). However, the frequency of these elongation events increased strongly with external force and elongation events were rarely observed at forces as low as 8pN (35) in agreement with the present study.

### Putative biological functions of the retraction ATPase

Recent reports (1, 2, 20) together with our current study show that T4P systems employing retraction ATPases tend to generate higher force and retract at higher speed compared to systems lacking retraction ATPases. T4P systems lacking retraction ATPases are associated with protein secretion (18), surface sensing (19), and DNA in uptake systems in gram positive bacteria (34). We propose that high force generation may not be required for these functions. DNA uptake in gram negative species is strongly impaired in the absence of the retraction ATPases (33, 43), suggesting a role of high force generation during threading into the periplasm. Twitching motility is another function of T4P (4). T4P elongate, adhere to a surface, and subsequently pull the cell body forward by retraction. The rupture forces of T4P from abiotic surfaces (5, 39) and from other T4P (17) are in the range of 50 pN. During cellular movement, T4P must detach from the surface, otherwise movement stalls (5, 46). Therefore, the motor force must exceed 50 pN consistent with PilT-powered retraction. Similarly, T4P retraction regulates the kinetics of T4P-T4P attachment and detachment within bacterial colonies (17). Colonies formed by *N. gonorrhoeae* and *N. meningiditis* generating functional PilT behave like liquids whereas *pilT*-deletion strains behave glass like (16, 17). Liquidlike behavior facilitates colonization of blood vessels during meningococcal infection (16) and cell sorting with respect to differential T4P-T4P adhesion forces (14). Another function of T4P retraction that requires high forces is signaling to host cells. When epithelial cells are infected with gonococci or mock-infected with T4P-coated beads, cytoskeletal proteins accumulate to the site of infection (8–10, 41). This accumulation depends on PilT and force application for gonococcal and mock-infection, respectively (9, 10). We conclude that T4P retraction in the absence of a retraction ATPase is sufficient for some T4P-related functions, but for those functions that call for high forces, retraction ATPases are required.

## CONCLUSION

We have shown that gonococcal T4P retract in the absence of the retraction ATPase PilT. Both the speed and the maximum force of *pilT*-independent retraction are by orders of magnitude lower compared to PilT-powered retraction, explaining why *pilT*-independent T4P retraction has been overlooked so far. Our findings together with recent results for other species strongly suggest that T4P retraction without PilT is a general phenomenon. We thus propose that the T4P elongation ATPase is necessary to provide energy for fibre formation while retraction occurs spontaneously. Figuring out the chemo-mechanical coupling within the T4P machine and especially the dynamics of the platform complex in the presence and absence of the retraction ATPase will be a future challenge.

## MATERIALS AND METHODS

### Growth conditions

Gonococcal base agar was made from 10 g/l BactoTM agar (BD Biosciences, Bedford, MA, USA), 5 g/l NaCl (Roth, Darmstadt, Germany), 4 g/l K2HPO4 (Roth), 1 g/l KH2PO4 (Roth), 15 g/l BactoTM Proteose Peptone No. 3 (BD), 0.5 g/l soluble starch (Sigma-Aldrich, St. Louis, MO, USA)) and supplemented with 1% IsoVitaleX: 1 g/l D-Glucose (Roth), 0.1 g/l L-glutamine (Roth), 0.289 g/l L-cysteine-HCL×H20 (Roth), 1 mg/l thiamine pyrophosphate (Sigma-Aldrich), 0.2 mg/l Fe (NO3)3 (Sigma-Aldrich), 0.03 mg/lthiamine HCl (Roth), 0.13 mg/l 4-aminobenzoic acid (Sigma-Aldrich), 2.5 mg/l β-nicotinamide adenine dinucleotide (Roth) and 0.1 mg/l vitamin B12 (Sigma-Aldrich). GC medium is identical to the base agar composition, but lacks agar and starch.

### Bacterial strains

All bacterial strains were derived from the gonococcal strain MS11 (VD300). In all strains, we deleted the G4-motif by replacing it with the *aac* gene conferring resistance against apramycin. The G4-motif is essential for antigenic variation of the major pilin subunit (47). Pilin antigenic variation modifies the primary sequence of the pilin gene. To generate strain *ΔpilT2 ΔpilT* (Ng184), Ng150 (48) was transformed with genomic DNA from *ΔpilT2*(36) and subsequently with DNA from strain *ΔpilT* (Ng178). To generate strain *ΔpilU ΔpilT* (Ng185), Ng150 (48) was transformed with genomic DNA from GU2 (49) and subsequently with DNA from strain *ΔpilT* (Ng178). To generate strain *ΔpilT2 ΔpilU ΔpilT* (Ng186), Ng150 (48) was transformed with genomic DNA from GU2 (49), *ΔpilT2* (36), and subsequently with DNA from strain *ΔpilT* (Ng178).

*3xpilE* (Ng188) carries two tandemly arrayed (identical) *pilE* genes expressed ectopically in addition to the *pilE* gene in the native locus (39). It was constructed essentially as described for strain GE21 (40) with the exception that it carries two gene copies in addition to the native copy resulting in a strain expressing three identical *pilE* genes. This strain was generated by transforming strain Ng150 (48) with genomic DNA from strain Ng088 (40) and subsequently with DNA from strain *ΔpilT* (Ng178).

Transformants were selected on agar plates containing the respective antibiotics (Table 1).

### Characterization of T4P retraction

Retraction velocities and stalling forces were measure during an optical tweezers setup assembled on a Zeiss Axiovert 200 (35). In short, all measurements were carried out in retraction assay medium consisting of phenol red-free Dulbecco’s modified Eagle’s medium (Gibco, Grand Island, NY) with 2 mM L-glutamine (Gibco), 8 mM sodium pyruvate (Gibco) and 30mM HEPES (Roth). A Suspension of Bacteria and carboxylated latex beads with a diameter of 2 μm (Polysciences, Warrington, PA) was applied to a microscope slide and sealed. All measurements were performed at 37 °C. The trap stiffness was determined by power spectrum analysis of the Brownian motion of a trapped bead to be 0.5 pN/nm ± 10%. The retraction velocities were measured in force clamp mode. During the experiment, a bead was trapped and held close to an immobilized bacterium at the surface. Eventually, a pilus attached to the bead and its retraction lead to a deflection out of the equilibrium position. As soon as the deflection of the bead reached the threshold deflection corresponding to a force of 4 or 8 pN, a force feedback algorithm held the displacement constant by moving the sample in the xy-plane using a piezo stage. Stalling forces were measured in position clamp mode.

### Double laser trap

In order to investigate single cell interactions, we followed a previously developed protocol (17). Two optical traps were installed in an inverted microscope (Nikon TE2000 C1). The trapping laser (20I-BL-106C, Spectra Physics, 1064nm, 5.4W) was directed into a water-immersion objective (Nikon Plan Apochromate VC 60x N.A. 1.20). Manipulation of the laser was done with a two-axis acousto-optical deflector (DTD-274HD6 Collinear Deflector, Intra Action Corp., USA). Bacterial interaction was recorded with a CCD camera (sensicam qe, PCO, Kelheim, Germany). The optical trap was calibrated via the equi partition and drag force methods. At 100% laser power, the average stiffness is 0.11 (± 0.01) pN/nm. The linear regime extends up to 80pN.

### Depletion of proton motive force

Cells were incubated with 50 μM CCCP for 15 min prior to usage in dual laser tweezers. To check that 15 min are sufficient to affect cells, twitching motility of *pilT*-expressing *ΔG4* cells was checked by bright field microscopy. Consistent with previous results (38), cells showed low speed twitching motility after 15 min of treatment with 50 μM CCCP and high speed twitching motility without CCCP.

## ACKNOWLEDGEMENTS

We are grateful to Katrina Forest, Lisa Craig, and the members of the Maier lab for helpful discussions. This work was supported by the Deutsche Forschungsgemeinschaft through grant MA3898.

---

**FIG S1.**
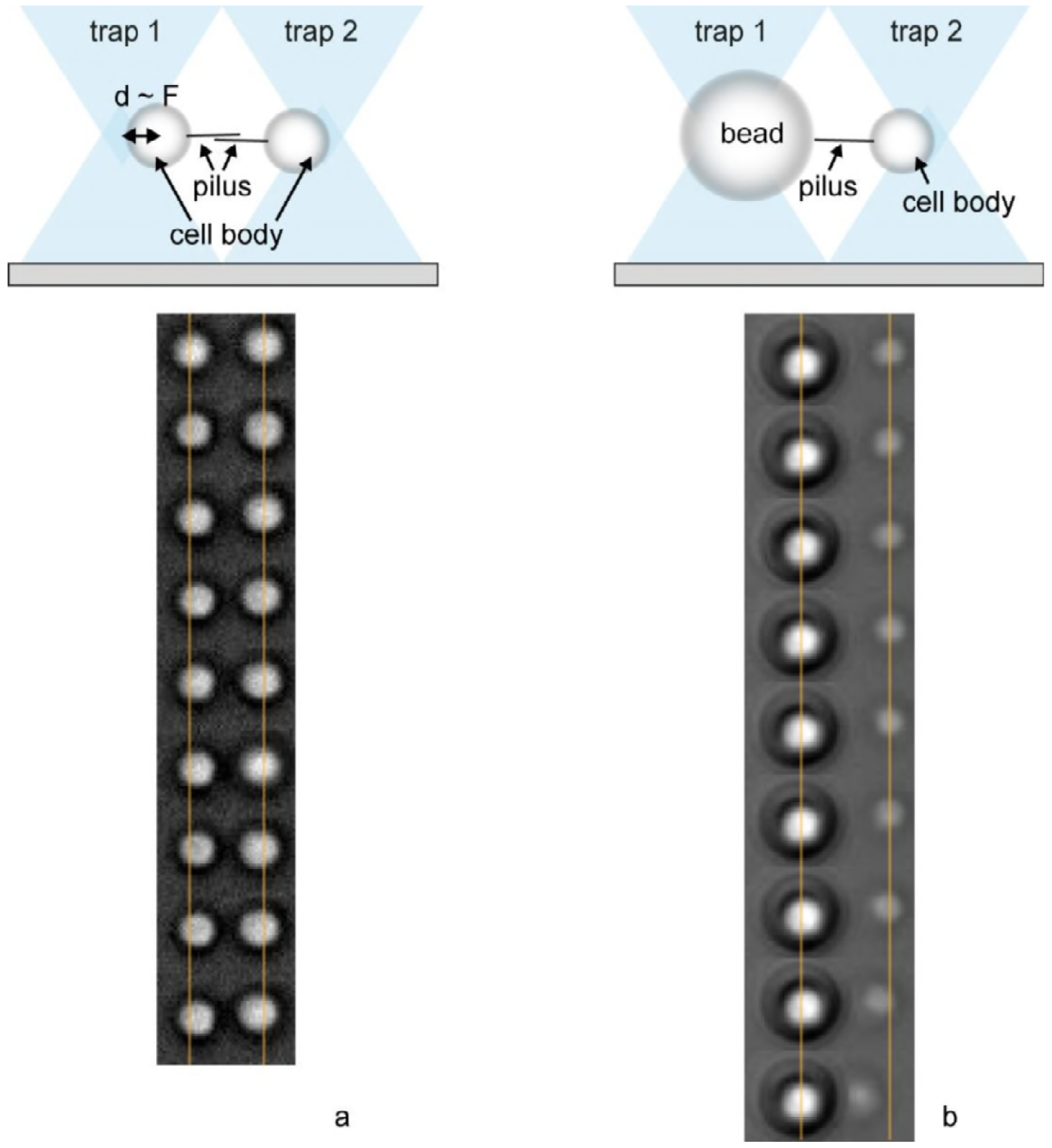
T4P retraction in *ΔpilT* strain (Ng178) in dual laser trap setup. a) A spherical gonococcus was trapped in each laser trap. Typical time lapse of two cells attracting each other. b) A bead was trapped in one trap and a spherical gonococcus in the other trap. Typical time lapse of a cells moving towards the bead. The laser trap exerts a stronger force on the bead than on the bacterium and therefore the deflection of the bacterium is higher. Δt = 6.53 s.

## References

1. Maier B, Potter L, So M, Long CD, Seifert HS, Sheetz MP. 2002. Single pilus motor forces exceed 100 pN. Proc Natl Acad Sci U S A 99:16012–16017.

2. Clausen M, Jakovljevic V, Sogaard-Andersen L, Maier B. 2009. High-force generation is a conserved property of type IV pilus systems. J Bacteriol 191:4633–4638.

3. Biais N, Ladoux B, Higashi D, So M, Sheetz M. 2008. Cooperative retraction of bundled type IV pili enables nanonewton force generation. PLoS Biol 6:e87.

4. Merz AJ, So M, Sheetz MP. 2000. Pilus retraction powers bacterial twitching motility. Nature 407:98–102.

5. Marathe R, Meel C, Schmidt NC, Dewenter L, Kurre R, Greune L, Schmidt MA, Muller MJ, Lipowsky R, Maier B, Klumpp S. 2014. Bacterial twitching motility is coordinated by a two dimensional tug-of-war with directional memory. Nat Commun 5:3759.

6. Zaburdaev V, Biais N, Schmiedeberg M, Eriksson J, Jonsson AB, Sheetz MP, Weitz DA. 2014. Uncovering the mechanism of trapping and cell orientation during Neisseria gonorrhoeae twitching motility. Biophys J 107:1523–1531.

7. Sabass B, Koch MD, Liu G, Stone HA, Shaevitz JW. 2017. Force generation by groups of migrating bacteria. Proc Natl Acad Sci U S A 114:7266–7271.

8. Merz AJ, Enns CA, So M. 1999. Type IV pili of pathogenic Neisseriae elicit cortical plaque formation in epithelial cells. Mol Microbiol 32:1316–1332.

9. Howie HL, Glogauer M, So M. 2005. The N. gonorrhoeae type IV pilus stimulates mechanosensitive pathways and cytoprotection through a pilT-dependent mechanism. PLoS Biol 3:e100.

10. Opitz D, Maier B. 2011. Rapid cytoskeletal response of epithelial cells to force generation by type IV pili. PLoS One 6:e17088.

11. Biais N, Higashi DL, Brujic J, So M, Sheetz MP. 2010. Force-dependent polymorphism in type IV pili reveals hidden epitopes. Proc Natl Acad Sci U S A 107:11358–11363.

12. Klausen M, Aaes-Jorgensen A, Molin S, Tolker-Nielsen T. 2003. Involvement of bacterial migration in the development of complex multicellular structures in Pseudomonas aeruginosa biofilms. Mol Microbiol 50:61–68.

13. Anyan ME, Amiri A, Harvey CW, Tierra G, Morales-Soto N, Driscoll CM, Alber MS, Shrout JD. 2014. Type IV pili interactions promote intercellular association and moderate swarming of Pseudomonas aeruginosa. Proc Natl Acad Sci U S A 111:18013–18018.

14. Oldewurtel ER, Kouzel N, Dewenter L, Henseler K, Maier B. 2015. Differential interaction forces govern bacterial sorting in early biofilms. Elife 4.

15. Hockenberry AM, Hutchens DM, Agellon A, So M. 2016. Attenuation of the Type IV Pilus Retraction Motor Influences Neisseria gonorrhoeae Social and Infection Behavior. MBio 7.

16. Bonazzi D, Lo Schiavo V, Machata S, Djafer-Cherif I, Nivoit P, Manriquez V, Tanimoto H, Husson J, Henry N, Chate H, Voituriez R, Dumenil G. 2018. Intermittent Pili-Mediated Forces Fluidize Neisseria meningitidis Aggregates Promoting Vascular Colonization. Cell 174:143–155 e116.

17. Welker A, Cronenberg T, Zollner R, Meel C, Siewering K, Bender N, Hennes M, Oldewurtel ER, Maier B. 2018. Molecular Motors Govern Liquidlike Ordering and Fusion Dynamics of Bacterial Colonies. Phys Rev Lett 121:118102.

18. Ng D, Harn T, Altindal T, Kolappan S, Marles JM, Lala R, Spielman I, Gao Y, Hauke CA, Kovacikova G, Verjee Z, Taylor RK, Biais N, Craig L. 2016. The Vibrio cholerae Minor Pilin TcpB Initiates Assembly and Retraction of the Toxin-Coregulated Pilus. PLoS Pathog 12:e1006109.

19. Ellison CK, Kan J, Dillard RS, Kysela DT, Ducret A, Berne C, Hampton CM, Ke Z, Wright ER, Biais N, Dalia AB, Brun YV. 2017. Obstruction of pilus retraction stimulates bacterial surface sensing. Science 358:535–538.

20. Ellison CK, Dalia TN, Vidal Ceballos A, Wang JC, Biais N, Brun YV, Dalia AB. 2018. Retraction of DNA-bound type IV competence pili initiates DNA uptake during natural transformation in Vibrio cholerae. Nat Microbiol 3:773–780.

21. Craig L, Volkmann N, Arvai AS, Pique ME, Yeager M, Egelman EH, Tainer JA. 2006. Type IV pilus structure by cryo-electron microscopy and crystallography: implications for pilus assembly and functions. Mol Cell 23:651–662.

22. Hospenthal MK, Costa TRD, Waksman G. 2017. A comprehensive guide to pilus biogenesis in Gram-negative bacteria. Nat Rev Microbiol 15:365–379.

23. Chang YW, Rettberg LA, Treuner-Lange A, Iwasa J, Sogaard-Andersen L, Jensen GJ. 2016. Architecture of the type IVa pilus machine. Science 351:aad2001.

24. McCallum M, Tammam S, Khan A, Burrows LL, Howell PL. 2017. The molecular mechanism of the type IVa pilus motors. Nat Commun 8:15091.

25. Kolappan S, Coureuil M, Yu X, Nassif X, Egelman EH, Craig L. 2016. Structure of the Neisseria meningitidis Type IV pilus. Nat Commun 7:13015.

26. Mancl JM, Black WP, Robinson H, Yang Z, Schubot FD. 2016. Crystal Structure of a Type IV Pilus Assembly ATPase: Insights into the Molecular Mechanism of PilB from Thermus thermophilus. Structure 24:1886–1897

27. Chang YW, Kjaer A, Ortega DR, Kovacikova G, Sutherland JA, Rettberg LA, Taylor RK, Jensen GJ. 2017. Architecture of the Vibrio cholerae toxin-coregulated pilus machine revealed by electron cryotomography. Nat Microbiol 2:16269.

28. Freitag NE, Seifert HS, Koomey M. 1995. Characterization of the pilF-pilD pilus-assembly locus of Neisseria gonorrhoeae. Mol Microbiol 16:575–586.

29. Wolfgang M, Lauer P, Park HS, Brossay L, Hebert J, Koomey M. 1998. PilT mutations lead to simultaneous defects in competence for natural transformation and twitching motility in piliated Neisseria gonorrhoeae. Mol Microbiol 29:321–330.

30. Wolfgang M, van Putten JP, Hayes SF, Dorward D, Koomey M. 2000. Components and dynamics of fiber formation define a ubiquitous biogenesis pathway for bacterial pili. EMBO J19:6408–6418.

31. Van Satyshur KA, Worzalla GA, Meyer LS, Heiniger EK, Aukema KG, Misic AM, Forest KT. 2007. Crystal structures of the pilus retraction motor PilT suggest large domain movements and subunit cooperation drive motility. Structure 15:363–376.

32. Misic AM, Satyshur KA, Forest KT. 2010. P. aeruginosa PilT structures with and without nucleotide reveal a dynamic type IV pilus retraction motor. J Mol Biol 400:1011–1021.

33. Seitz P, Blokesch M. 2013. DNA-uptake machinery of naturally competent Vibrio cholerae. Proc Natl Acad Sci U S A 110:17987–17992.

34. Chen I, Dubnau D. 2004. DNA uptake during bacterial transformation. Nat Rev Microbiol l2:241–249.

35. Clausen M, Koomey M, Maier B. 2009. Dynamics of type IV pili is controlled by switching between multiple states. Biophys J 96:1169–1177.

36. Kurre R, Hone A, Clausen M, Meel C, Maier B. 2012. PilT2 enhances the speed of gonococcal type IV pilus retraction and of twitching motility. Mol Microbiol 86:857–865.

37. Kurre R, Maier B. 2012. Oxygen depletion triggers switching between discrete speed modes of gonococcal type IV pili. Biophys J 102:2556–2563.

38. Kurre R, Kouzel N, Ramakrishnan K, Oldewurtel ER, Maier B. 2013. Speed switching of gonococcal surface motility correlates with proton motive force. PLoS One 8:e67718.

39. Holz C, Opitz D, Greune L, Kurre R, Koomey M, Schmidt MA, Maier B. 2010. Multiple pilus motors cooperate for persistent bacterial movement in two dimensions. Phys Rev Lett 104:178104.

40. Aas FE, Winther-Larsen HC, Wolfgang M, Frye S, Lovold C, Roos N, van Putten JP, Koomey M. 2007. Substitutions in the N-terminal alpha helical spine of Neisseria gonorrhoeae pilin affect Type IV pilus assembly, dynamics and associated functions. Mol Microbiol 63:69–85.

41. Higashi DL, Lee SW, Snyder A, Weyand NJ, Bakke A, So M. 2007. Dynamics of Neisseria gonorrhoeae attachment: microcolony development, cortical plaque formation, and cytoprotection. Infect Immun 75:4743–4753.

42. Hepp C, Maier B. 2016. Kinetics of DNA uptake during transformation provide evidence for a translocation ratchet mechanism. Proc Natl Acad Sci U S A 113:12467–12472.

43. Gangel H, Hepp C, Muller S, Oldewurtel ER, Aas FE, Koomey M, Maier B. 2014. Concerted spatio-temporal dynamics of imported DNA and ComE DNA uptake protein during gonococcal transformation. PLoS Pathog 10:e1004043.

44. Ribbe J, Baker AE, Euler S, O’Toole GA, Maier B. 2017. Role of Cyclic Di-GMP and Exo polysaccharide in Type IV Pilus Dynamics. J Bacteriol 199.

45. Maier B, Koomey M, Sheetz MP. 2004. A force-dependent switch reverses type IV pilus retraction. Proc Natl Acad Sci U S A 101:10961–10966.

46. Ponisch W, Weber CA, Juckeland G, Biais N, Zaburdaev V. 2017. Multiscale modeling of bacterial colonies: how pili mediate the dynamics of single cells and cellular aggregates. New Journal of Physics 19.

47. Rotman E, Seifert HS. 2014. The genetics of Neisseria species. Annu Rev Genet 48:405–431.

48. Zollner R, Oldewurtel ER, Kouzel N, Maier B. 2017. Phase and antigenic variation govern competition dynamics through positioning in bacterial colonies. Scientific Reports 7.

49. Park HS, Wolfgang M, Koomey M. 2002. Modification of type IV pilus-associated epithelial cell adherence and multicellular behavior by the PilU protein of Neisseria gonorrhoeae. Infect Immun 70:3891–3903.

